# Inferring node dates from tip dates in fossil Canidae: the importance of tree priors

**DOI:** 10.1101/049643

**Authors:** Nicholas J. Matzke, April Wright

## Abstract

Tip-dating methods are becoming popular alternatives to traditional node calibration approaches for building time-scaled phylogenetic trees, but questions remain about their application to empirical datasets. We compared the performance of the most popular methods against a dated tree of fossil Canidae derived from previously published monographs. Using a canid morphology dataset, we performed tip-dating using Beast 2.1.3 and MrBayes 3.2.5. We find that for key nodes (*Canis*, ~3.2 Ma, Caninae ~11.7 Ma) a non-mechanistic model using a uniform tree prior produces estimates that are unrealistically old (27.5, 38.9 Ma). Mechanistic models (incorporating lineage birth, death, and sampling rates) estimate ages that are closely in line with prior research. We provide a discussion of these two families of models (mechanistic vs. non-mechanistic) and their applicability to fossil datasets.

## Main text

“Tip-dating” methods allow for fossils to be incorporated as terminal taxa in divergence dating analysis. These methods require a tree model that allows non-contemporaneous tips. These models can be categorized broadly into two types: mechanistic models where trees are a function of parameterized speciation, extinction, and sampling processes, termed birth-death-serial-sampling (BDSS; [2]) or fossilized birth-death (FBD; [3]) models; and the non-mechanistic uniform prior on trees and node ages [1], which does not have parameters for the rates of these processes. BDSS/FBD models can allow or disallow sampled ancestors (SA; [4, 5]). Importantly, tip-dating methods allow researchers to avoid relying on node calibrations. While node calibration approaches are valuable, they are subject to a number of well-known criticisms [1, 3, 6–8] such as subjectivity and incomplete use of information. Node calibration also weakens inferential capacity by requiring *a priori* constraint of dates that researchers would prefer to infer.

As a result of these analytical advantages, tip-dating methods are becoming popular. However, some studies using these approaches on empirical datasets have reached negative conclusions about the plausibility of inferred dates (references in Supplemental Material, SM). While tip-dating methods have been validated against simulations, it is debatable to what extent manufactured histories are comparable to the complexity of real evolutionary histories [9]. For empirical work, it can be difficult to tell if problematic inferences in a particular study are due to the data, the methods, human error or a combination of the three.

It may therefore be useful to compare tip-dating inferences on a high-quality empirical dataset, one where the fossil record strongly corroborates key divergence times without Bayesian computational methods. An ideal dataset would also avoid difficulties found in classic dating questions such as the origin of angiosperms, placental mammals, crown birds, and the Cambrian phyla (Table 1). Suitable fossil datasets are rare, but one for which a strong argument (Table 1) can be made is the fossil Canidae (dog family; [10]). Monographs on the three Canidae subfamilies Hesperocyoninae [11], Borophaginae [12], and Caninae [13] combined cladistic analysis of discrete characters with expert knowledge of stratigraphy and continuous characters to produce species-level phylogenies dated to ~1-2 my resolution. We use Canidae to compare date estimates made under mechanistic (BDSS/FBD) and non-mechanistic (uniform tree prior) models to expert opinion. We conclude that reasonable date estimation requires an appropriate choice of tree prior, which may vary by paleontological dataset.

**Table 1.**
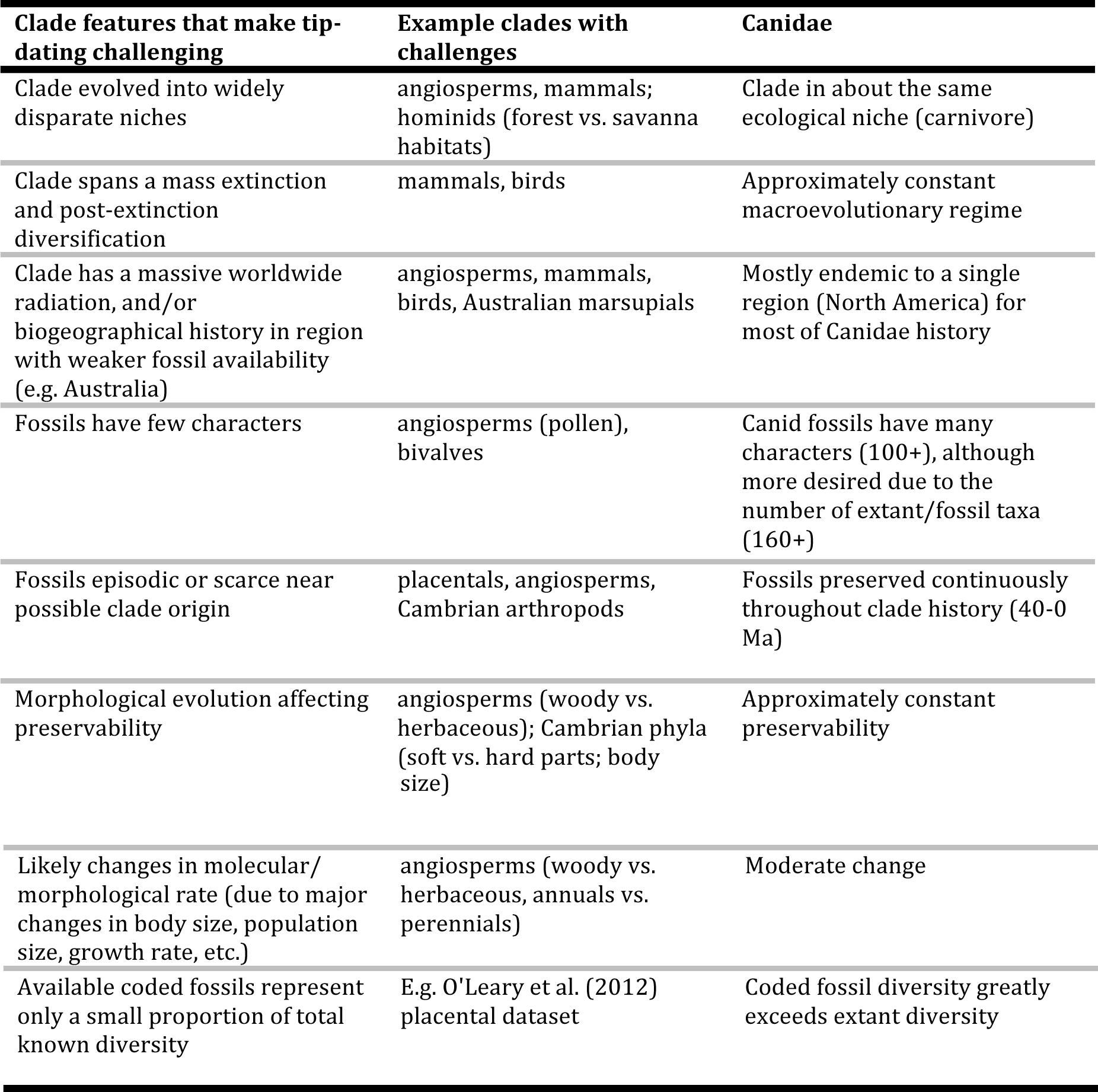
Clade features that present challenges to tip-dating methods (or any dating methods). Canidae exhibit few of the issues that may confound dating in other clades (e.g. angiosperms, mammals, birds).

## Methods

### Data

The “expert tree” was digitized from the monographs of Wang and Tedford [11–13] using TreeRogue [14], with judgment calls resolved in favour of preserving the authors’ depiction of divergence times (SM). Morphological characters and dates came from Slater (2015) [15, 16].

### Tip-dating analyses

MrBayes analyses were conducted by modification of Slater’s commands file. 58 variants of MrBayes analyses were constructed to investigate several issues noticed in the interaction of MrBayes versions and documentation, and Slater’s commands file (SM, Appendix 1).

We compared the expert tree (Figure 1a) and Slater’s published uniform tree prior analysis which included many node-date constraints (Figure 1b: mb1_orig) to six focal analyses (four MrBayes3.2.5 analyses and two Beast2.1.3). These were (1c) *mb1_UC*: Slater’s analysis with various corrections; (1d) *mb8_UU*: uniform tree prior, uninformative priors on clock parameters, and no node date calibrations except for a required root age calibration, set to uniform(45,100) to represent the common situation where researchers wish to infer node dates rather than pre-specify them; (1f) *mb9x_SA*: mb8_UU but with SA-BDSS tree prior and flat priors on speciation, extinction, and sampling rate; (1e) *mb10_noSA*: mb9x_SA but noSA-BDSS, i.e. disallowing sampled ancestors; (1g) *r1_noSA*: Beast2 noSA-BDSS analysis with flat priors used for each major parameter (mean and SD of the lognormal relaxed clock; and birth, death, and serial sampling rates); (1h) *r2_SA*: Beast2 SA-BDSS analysis with the same priors. Beast2 analyses were constructed with BEASTmasteR [17, 18]; full details on the analyses are in SM.

**Figure 1.**
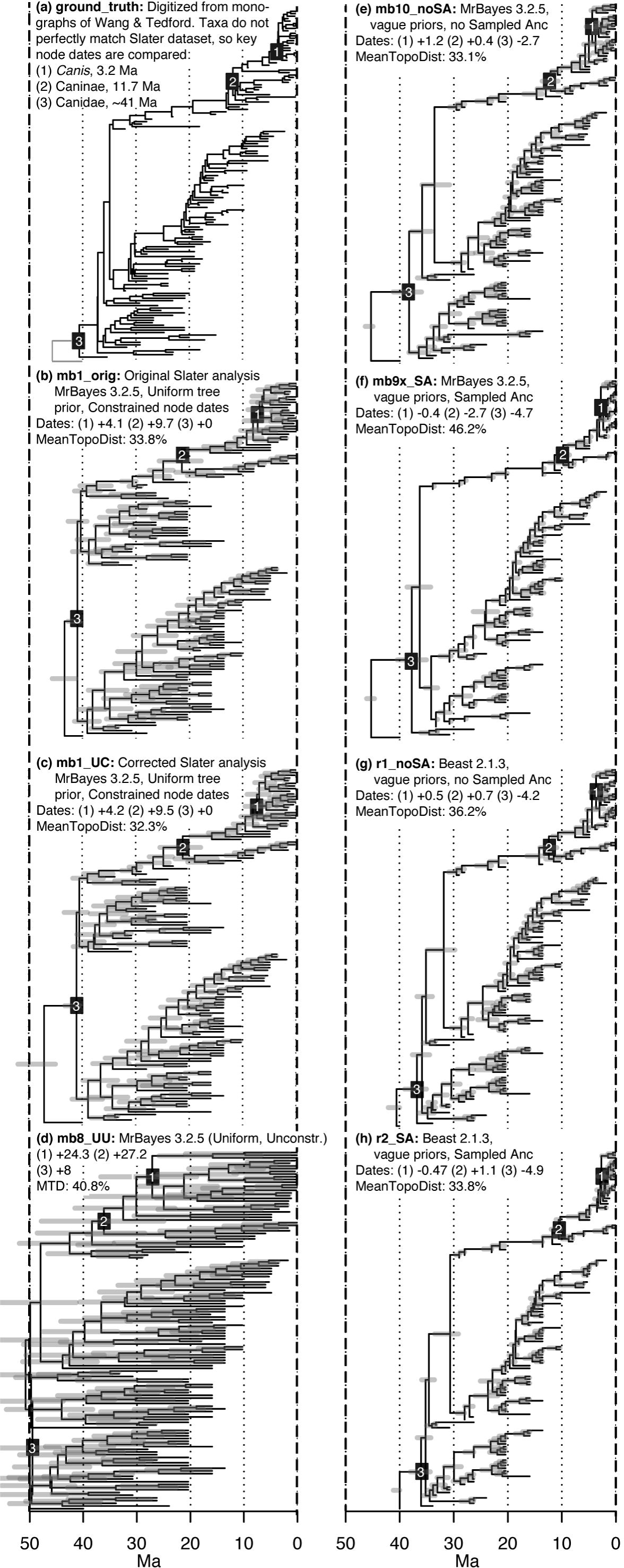
Comparison of (a) the expert tree, to seven Bayesian dating analyses (b-h) using the Slater (2015) characters and dates. As the expert tree’s taxa do not perfectly overlap with the Slater taxa, key node dates are compared: (1) the common ancestor of crown (extant) *Canis*, (2) the common ancestor of living Caninae, and (3) the common ancestor of the total group Canidae. The expert tree dates are given in (a), and the differences from these are given in (b-h). The percentages represent the Mean Topological (RF) Distances between (b-h) and mb2_undated (average within-posterior distance=24.6%). Note: The expert tree lacks Slater’s “outgroup” OTU (the branch below node 3), but this has been added for visual comparability (grey line).

## Results

The six focal analyses are compared in Figure 1, and key priors and results are shown in Table S1. The unconstrained MrBayes uniform tree prior analysis (mb8_UU) produces estimates with implausibly old ages and huge uncertainties, and with the age of Canidae overlapping the K-Pg boundary. This behaviour was also noted by Slater [15]. The expert-tree dates of crown *Canis* (which includes *Cuon*, *Lycaon*, and *Xenocyon*) and crown Caninae are ~3.2 and ~11.7 Ma, but mb8_UU makes mean estimates of 27.5 and 38.9 Ma, and even the wide 95% highest posterior densities (HPDs), spanning 22-25 my, do not overlap expert opinion. More surprisingly, even Slater’s highly constrained analysis (mb1_UC), although closer, does not produce HPDs (5.1-9.6 Ma; 17.8-25.5 Ma) that overlap expert-tree dates. In contrast, both Beast2 estimates (r1_noSA and r2_SA) and MrBayes noSA-BDSS (mb10_noSA, mb9x_SA) are within ~1-2 Ma of expert estimates, HPD widths ~2-3 my). The date of total-group Canidae (node 3, Figure 1) matches the expert tree when it has been constrained (mb1_UC), but is 27 Ma older in mb8_UU, and consistently ~3-5 Ma younger in BDSS-type analyses.

Additional comparisons are available in SM and Tables S1-S2, including comparisons of topological distances between the Bayesian dating estimates and an undated MrBayes analysis on the same data and posterior prediction of tip dates. The SM and Appendix 1 also discuss difficulties observed in some non-focal runs.

## Discussion

The result of greatest interest is the contrast between expert-tree dates and dates inferred with the uniform tree prior. Whether or not this is surprising may depend on researcher background. We suggest that reasoning from first principles suggests that effective tip-dating under the uniform tree prior will be difficult without strongly informative priors on node dates and/or clock rate and variability. Apart from such constraints, nothing in the tip dates or the uniform tree prior restricts the age of nodes below the dated tips; thus, in our fossils-only analysis, the node ages are scaled up and down as the root age is sampled according to the root age prior. Without informative priors, the clock rate and variability parameters will adjust along with the tree height; highly uncertain node ages will result.

Despite what first principles suggest, we suspect our results may surprise some researchers. The MrBayes uniform tree prior was the leading model in the early tip-dating literature (11/16 papers as of mid-2015, 9 of them as the exclusive Bayesian tip-dating method; SM), and until recently (October 2014, v. 3.2.3) the uniform tree prior was the only option available in MrBayes. Early tip-dating efforts in Beast/Beast2 required tedious manual editing of XML and/or elaborate scripting efforts (such as BEASTmasteR), whereas MrBayes was relatively easy to use. Therefore, many early attempts at tip-dating used the uniform tree prior.

In contrast to the results with the uniform tree prior, analyses using BDSS/FBD tree priors (mb10_noSA, mb9x_SA, r1_noSA, r2_SA) retrieved results that approximate previous age estimates. Given only the characters and tip-dates, and with uninformative priors on parameters and the root age, these analyses were able to estimate node ages that were close to expert opinion, with a high rate of fossil sampling limiting node ages. These analyses gave more reasonable age and uncertainty estimates than the uniform tree prior even when the uniform was given substantial additional information in the form of many node calibrations (mb1_UC). Even well constrained uniform tree prior analyses displayed a tendency to space node ages evenly between calibrations and tip dates, regardless of morphological branch lengths (SM).

Tip-dating with the uniform tree prior was introduced [1] as an alternative to node calibration, attractive because tip-dating avoided various undesirable compromises that researchers are forced to make to when constructing node-age priors. Ronquist et al. [1] also critiqued Stadler’s [2] BDSS prior as being “complete but unrealistic,” particularly due to assumptions about constant birth/death/sampling rates and sampling in the Recent. They offered the uniform prior as an alternative, free of these difficulties. If, however, strongly informative priors on rates or node age calibrations are required to produce reasonable results under the uniform tree prior, its main appeal is lost. The addition of BDSS/FBD models with sampled ancestors to MrBayes [5] suggests that the best prospects for tip-dating may lay in adding realism to mechanistic models, rather than in attempting to devise nonmechanistic, agnostic dating priors.

A major caveat in our study is that we did not attempt to study the effect of poorer fossil taxon sampling on the inferences made under different tree priors. Canidae are unusually well sampled. In other cases researchers may only have a handful of fossils when true diversity was hundreds or thousands of species (closer to the situation in the exemplar Hymenoptera dataset explored by [1, 5]). In such situations the uniform tree prior’s performance may improve relative to BDSS-type models attempting to estimate mechanistic parameters from few data.

A great deal of work remains to understand how best to perform tip-dating analyses. We have shown that for this high-quality dataset, mechanistic and nonmechanistic models perform quite differently, and present an argument that mechanistic models are more appropriate for this dataset.

## Data accessibility

All scripts, data files, and results files are available via a zipfile on Dryad (doi: 10.5061/dryad.vn52f)

## Competing interests

We have no competing interests.

## Authors’ Contributions

NJM wrote *BEASTmasteR*, conducted the Beast2 computational analyses and drafted the manuscript. AW contributed to MrBayes dating efforts and edited and corrected the manuscript. Both authors agree to be held accountable for the content therein and approve the final version of the manuscript.

## Acknowledgements

We thank David Bapst, Graeme Lloyd, Jeremy Beaulieu, Kathryn Massana, Brian O’Meara, Graham Slater, and Mike Lee for helpful comments and discussion, as well as the participants of the 2014 Society of Vertebrate Paleontology tip-dating workshop/symposium. We also thank the BEAST developers and the *beast-users* Google Group, particularly Remco Bouckaert.

### Funding

NJM was supported by NIMBioS fellowship under NSF Award #EFJ0832858, and ARC DECRA fellowship DE150101773. Work on this topic began under the NSF Bivalves in Time and Space grant (DEB-0919451). AW was supported by NSF DEB-1256993.

## References

[1] Ronquist, F., Klopfstein, S., Vilhelmsen, L., Schulmeister, S., Murray, D.L. & Rasnitsyn, A.P. 2012 A total-evidence approach to dating with fossils, applied to the early radiation of the Hymenoptera. Systematic Biology 61, 973–999. (doi:10.1093/sysbio/sys058).

[2] Stadler, T. 2010 Sampling-through-time in birth-death trees. Journal of Theoretical Biology 267, 396–404. (doi:10.1016/j.jtbi.2010.09.010).

[3] Heath, T.A., Huelsenbeck, J.P. & Stadler, T. 2014 The fossilized birth-death process for coherent calibration of divergence-time estimates. Proceedings of the National Academy of Sciences 111, E2957–E2966. (doi:10.1073/pnas.1319091111).

[4] Gavryushkina, A., Welch, D., Stadler, T. & Drummond, A.J. 2014 Bayesian inference of sampled ancestor trees for epidemiology and fossil calibration. PLoS Comput Biol 10, e1003919. (doi:10.1371/journal.pcbi.1003919).

[5] Zhang, C., Stadler, T., Klopfstein, S., Heath, T.A. & Ronquist, F. 2016 Total-evidence dating under the fossilized birth-death process. Systematic Biology 65, 228–249. (doi:10.1093/sysbio/syv080).

[6] Parham, J.F., Donoghue, P.C.J., Bell, C.J., Calway, T.D., Head, J.J., Holroyd, P.A., Inoue, J.G., Irmis, R.B., Joyce, W.G., Ksepka, D.T., Patané, J.S.L., Smith, N.D., Tarver, J.E., van Tuinen, M., Yang, Z., Angielczyk, K.D., Greenwood, J.M., Hipsley, C.A., Jacobs, L., Makovicky, P.J., Müller, J., Smith, K.T., Theodor, J.M., Warnock, R.C.M. & Benton, M.J. 2012 Best practices for justifying fossil calibrations. Systematic Biology 61, 346–359. (doi:10.1093/sysbio/syr107).

[7] Pyron, R.A. 2011 Divergence time estimation using fossils as terminal taxa and the origins of Lissamphibia. Systematic Biology 60, 466–481. (doi:10.1093/sysbio/syr047).

[8] Wood, H.M., Matzke, N.J., Gillespie, R.G. & Griswold, C.E. 2013 Treating fossils as terminal taxa in divergence time estimation reveals ancient vicariance patterns in the Palpimanoid spiders. Systematic Biology 62, 264–284. (doi:10.1093/sysbio/sys092).

[9] Hillis, D.M. 1995 Approaches for assessing phylogenetic accuracy. Systematic Biology 44, 3–16. (doi:10.1093/sysbio/44.1.3).

[10] Wang, X.T., Richard H. 2008 Dogs: their fossil relatives and evolutionary history. New York, Columbia University Press.

[11] Wang, X. 1994 Phylogenetic systematics of the Hesperocyoninae (Carnivora, Canidae). Bulletin of the American Museum of Natural History 221, 1–207.

[12] Wang, X.T., Richard H.; Taylor, Beryl E. 1999 Phylogenetic systematics of the Borophaginae. Bulletin of the American Museum of Natural History 243, 1–391.

[13] Tedford, R.H.W., Xiaoming; Taylor, Beryl E. 2009 Phylogenetic systematics of the North American fossil Caninae (Carnivora, Canidae). Bulletin of the American Museum of Natural History 325, 1–218.

[14] Matzke, N.J. 2013 TreeRogue: R code for digitizing trees. https://stat.ethz.ch/pipermail/r-sig-phylo/2010-October/000816.html

[15] Slater, G.J. 2015 Iterative adaptive radiations of fossil canids show no evidence for diversity-dependent trait evolution. Proceedings of the National Academy of Sciences 112, 4897–4902. (doi:10.1073/pnas.1403666111).

[16] Slater, G.J. 2015. Data from: Iterative adaptive radiations of fossil canids show no evidence for diversity-dependent trait evolution. Dryad. Accessed May 1, 2015. http://dx.doi.org/10.5061/dryad.9qd51

[17] Matzke, N.J. 2015 BEASTmasteR: automated conversion of NEXUS data to BEAST2 XML format, for fossil tip-dating and other uses. PhyloWiki. http://phylo.wikidot.com/beastmaster

[18] Matzke, N.J. 2016 The evolution of antievolution policies after *Kitzmiller versus Dover*. Science 351, 28–30. (doi:10.1126/science.aad4057).

